# Avian influenza viruses in wild birds: virus evolution in a multi-host ecosystem

**DOI:** 10.1101/282491

**Authors:** Divya Venkatesh, Marjolein J. Poen, Theo M. Bestebroer, Rachel D. Scheuer, Oanh Vuong, Mzia Chkhaidze, Anna Machablishvili, Jimsher Mamuchadze, Levan Ninua, Nadia B. Fedorova, Rebecca A. Halpin, Xudong Lin, Amy Ransier, Timothy B Stockwell, David E. Wentworth, Divya Kriti, Jayeeta Dutta, Harm van Bakel, Anita Puranik, Marek J Slomka, Steve Essen, Ian H. Brown, Ron A.M. Fouchier, Nicola S. Lewis

## Abstract

Wild ducks and gulls are the major reservoirs for avian influenza A viruses (AIVs). The mechanisms that drive AIV evolution are complex at sites where various duck and gull species from multiple flyways breed, winter or stage. The Republic of Georgia is located at the intersection of three migratory flyways: Central Asian Flyway, East Asian/East African Flyway and Black Sea/Mediterranean Flyway. For six consecutive years (2010-2016), we collected AIV samples from various duck and gull species that breed, migrate and overwinter in Georgia. We found substantial subtype diversity of viruses that varied in prevalence from year to year. Low pathogenic (LP)AIV subtypes included H1N1, H2N3, H2N5, H2N7, H3N8, H4N2, H6N2, H7N3, H7N7, H9N1, H9N3, H10N4, H10N7, H11N1, H13N2, H13N6, H13N8, H16N3, plus two H5N5 and H5N8 highly pathogenic (HP)AIVs belonging to clade 2.3.4.4. Whole genome phylogenetic trees showed significant host species lineage restriction for nearly all gene segments and significant differences for LPAIVs among different host species in observed reassortment rates, as defined by quantification of phylogenetic incongruence, and in nucleotide diversity. Hemagglutinin clade 2.3.4.4 H5N8 viruses, circulated in Eurasia during 2014-2015 did not reassort, but analysis after its subsequent dissemination during 2016-2017 revealed reassortment in all gene segments except NP and NS. Some virus lineages appeared to be unrelated to AIVs in wild bird populations in other regions with maintenance of local AIV viruses in Georgia, whereas other lineages showed considerable genetic inter-relationship with viruses circulating in other parts of Eurasia and Africa, despite relative under-sampling in the area.

**Importance:** Waterbirds (e.g., gulls/ducks) are natural reservoirs of avian influenza viruses (AIVs) and have been shown to mediate dispersal of AIV at inter-continental scales during seasonal migration. The segmented genome of influenza viruses enables viral RNA from different lineages to mix or re-assort when two viruses infect the same host. Such reassortant viruses have been identified in most major human influenza pandemics and several poultry outbreaks. Despite their importance, we have only recently begun to understand AIV evolution and reassortment in their natural host reservoirs. This comprehensive study illustrates of AIV evolutionary dynamics within a multi-host ecosystem at a stop-over site where three major migratory flyways intersect. Our analysis of this ecosystem over a six-year period provides a snapshot of how these viruses are linked to global AIV populations. Understanding the evolution of AIVs in the natural host is imperative to both mitigating the risk of incursion into domestic poultry and potential risk to mammalian hosts including humans.

## Introduction

Avian influenza viruses (AIVs) have been identified in a wide diversity of wild and domestic bird species but wild waterbirds of the Orders *Anseriformes and Charadriformes*, such as ducks, geese, swans and shorebirds (1, 2) form their natural reservoir. These birds maintain diverse group of low pathogenic avian influenza A viruses (LPAIVs), which cause limited morbidity in these host species in experimental settings (3). The effect of AIV infection in wild birds in non-experimental settings is more contradictory. Body mass was significantly lower in infected mallards (*Anas playrhynchos)* and the amount of virus shed by infected juveniles was negatively correlated with body mass. However, there was no general effect of infection on staging time (duration of stopover for migratory birds), except for juveniles in September and LPAIV infection did not affect speed or distance of subsequent migration (4). Conversely, a recent mallard study demonstrated no obvious detriment to the bird as movement patterns did not differ between LPAIV infected and uninfected birds. Hence, LPAIV infection probably does not affect mallard movements during stopover, consequently resulting in the potential for virus spread along the migration route (5). The precise role of migrants and resident birds in amplifying and dispersing AIVs however, remains unclear. In another study the migrant arrivals played a role in virus amplification rather than seeding a novel variant into a resident population (6). It has also been suggested that switching transmission dynamics might be a critical strategy for pathogens such as influenza A viruses associated with mobile hosts such as wild waterbirds, and that both intra and inter-species transmission are important to maintaining gene flow across seasons (7).

AIVs continue to cause both morbidity and mortality in poultry worldwide. Increased mortality is strongly related to infection with highly pathogenic influenza A viruses (HPAIVs), characterised by mortality in gallinaceous poultry (8). Periodically, human infections associated with HPAIV of both the H5 and H7 subtypes have been detected. In particular, parts of Asia and Africa have been significantly affected by the Eurasian (goose/Guangdong/1996) lineage H5 HPAIV epizootic for two decades, becoming enzootic in some areas and multiple waves of influenza with evolving viruses in others (9). More recently, H5Nx reassortants of the Eurasian lineage HPAIVs from clade have been introduced into wild birds from poultry and spread to new geographic regions (10).

The Caucasus, at the border of Europe and Asia, is important for migration and over-wintering of wild waterbirds. Three flyways, the Central Asian, East Africa-West Asia, and Mediterranean/Black Sea flyways, converge in this region. Understanding the ecology and evolution of AIVs in wild birds is complex, particularly at sites where multiple species co-habit and in those ecosystems which support different annual life-cycle stages and where multiple migratory flyways intersect.

At a population level, Eurasian dabbling ducks were found to be more frequently infected than other ducks and Anseriformes (11) with most AIV subtypes detected in ducks, except H13 and H16 subtypes which were detected primarily in gulls (11, 12). Temporal and spatial variation in influenza virus prevalence in wild birds was observed, with AIV prevalence varying by sampling location. In this study site in the Republic of Georgia, we observed peak prevalence in large gulls during the autumn migration (5.3-9.8%), but peak prevalence in Black-headed Gulls (*Chroicocephalus ridibundus*) in spring (4.2-13%)(13). In ducks, we observed increased AIV prevalence during the autumn post-moult aggregations and migration stop-over period (6.3%) but at lower levels to those observed in other more northerly post-moult areas in Eurasia.

In North America, studies have primarily focused on Anseriformes species with sampling during late summer and autumn southern migration (14-16), rather than longitudinally throughout the annual lifecycle of the host or within an ecosystem. The southwestern Lake Erie Basin is an important stopover site for waterfowl during migration periods, and over the past 28 years, 8.72% of waterfowl sampled in this geographic location have been positive for AIV recovery during summer and autumn (June – December) (17). More recent studies which targeted overwintering and returning migratory birds during February – April showed the presence of diverse AIV subtypes in waterbirds at northern latitudes in the United States (17).

Previous genetic studies of the viruses isolated from wild birds have focused on gene flow at an intra- or intercontinental level involving multiple hosts, rather than on virus gene flow among species within an ecosystem (16, 18-20). Indeed, the conclusions of such studies have been somewhat limited at times by statistical power owing to insufficient sequence data from enough hosts relevant to virus dynamics across the geographic study area. (21). In Eurasia, frequent reassortment and co-circulating lineages were observed for all eight genomic RNA segments over time. Although, there was no apparent species-specific effect on the diversity of the AIVs, there was a spatial and temporal relationship between the Eurasian sequences and significant viral migration of AIVs from West Eurasia towards Central Eurasia (22).

This study presents novel findings concerning the ecology and evolution of both LPAIVs and HPAIVs circulating in wild birds in a key active surveillance site in Eurasia. We investigated the diffusion of AIV gene segments within different wild bird hosts occupying the same ecosystem. There was substantial diversity in surface glycoprotein HA (heamagglutinin) and NA (neuraminidase) subtypes, which varied year to year and with the host species. M, NS, NP, PB1, PB2 and PA (henceforth referred to as “internal” gene segments) also showed host restriction to various degrees. There were differences in genetic diversity, reassortment rates, and inter-species transmission rates in the internal gene segments associated with different host species and HA subtypes. We also examined how closely related the Georgian AIV gene segments were to AIV globally. We found evidence for genetic inter-relationship of Georgian AIV with AIV in mainly Africa and Eurasia but several lineages appear to be maintained locally.

## Methods

Active surveillance for influenza A viruses was carried out from 2010-2016 as described previously (13)

### Dataset and genomic sequencing

Over a period of six years, 30,911 samples from 105 different bird species were analysed for the presence of AIVs. Positive isolates were obtained by standard approaches (23), and where possible, subtyped and sequence generated from extracted RNA as described below.

For virus samples from 2010-2012, codon complete genomes of IAV were sequenced as part of the Influenza Genome Project (http://gcid.jcvi.org/projects/gsc/influenza/index.php), an initiative by the National Institute of Allergies and Infectious Diseases (NIAID). IAV viral RNA (vRNA) was isolated from the samples/specimens, and the entire genome was amplified from 3 ul of RNA template using a multi-segment RT-PCR strategy (M-RTPCR) (24, 25). The amplicons were sequenced using the Ion Torrent PGM (Thermo Fisher Scientific, Waltham, Massachusetts, USA) and/or the Illumina MiSeq v2 (Illumina, Inc., San Diego, California, USA) instruments. When sequencing data from both platforms was available, the data were merged and assembled together; the resulting consensus sequences were supported by reads from both technologies. Sequence data for Georgia was downloaded from the NIAID Influenza Research Database (IRD) (Squires et al. 2012) through the web site at http://www.fludb.org on 11/5/2016. To this dataset, we added sequence data for isolates from 2013 and 2016 which were sequenced at either Erasmus MC, Animal and Plant Health Agency (APHA) or the Icahn School of Medicine at Mount Sinai (ISMMS). At Erasmus MC sequencing was performed as described previously by V. J. Munster et al. (26), with modifications. Primer sequences are available upon request.

At APHA, viral RNA was extracted using the QIAquick Viral RNA extraction kit (Qiagen, UK) without the addition of carrier. Double stranded cDNA (cDNA synthesis system, Roche, UK) was generated from RNA according to the manufacturer’s instructions. This was quantified using the fluorescent PicoGreen reagent and 1ng was used as a template for the preparation of the sequencing library (NexteraXT, Illumina, Cambridge, UK). Sequencing libraries were run on a MiSeq instrument (Illumina, Cambridge, UK) with 2×75 base paired end reads. Data handling of raw sequence reads and extraction of consensus sequences were performed at APHA.

For the Icahn School of medicine at Mount Sinai, RNA was extracted using the QIAamp Viral RNA Mini Kit (52904, Qiagen, UK). MS-RTPCR amplification was performed with the Superscript III high-fidelity RT-PCR kit (12574-023, Invitrogen) according to manufacturer’s instructions using the Opti1 primer set: Opti1-F1 5’ GTTACGCGCC<u>AGCAAAAGCAGG</u>, Opti1-F2 5’GTTACGCGCC<u>AGC**G**AAAGCAGG and</u> Opti1-R15’GTTACGCGCC<u>AGTAGAAACAAGG. DNA amplicons were purified using Agencourt AMPure XP 5ml Kit (A63880, Beckman Coulter). At the Icahn School of Medicine, sequencing libraries were prepared and sequencing was performed on a MiSeq instrument (Illumina, Cambridge, UK) with 2×150 base paired end reads.</u> Data handling of raw sequence reads and extraction of consensus sequences were performed at ISMMS, as described previously (27).

### Genetic analyses

#### Sequence alignment preparation

Whole genome sequences from 81 Georgian strains isolated between 2010 and 2016 are used in this analysis. We aligned sequences from each gene segment separately using MAFFT v7.305b (28) and trimmed to starting ATG and STOP codon in Aliview v1.18. Hemagglutinin (HA) sequences were further trimmed to exclude the initial signal sequence (29, 30). Sequences were then aligned using “muscle-codon” option with default settings in MEGA7 (31).

The NS gene has two alleles A and B, with significant difference in sequence composition, which could skew analyses of sequence diversity. The NS gene sequences were therefore considered both as a complete dataset (NS) and subdivided into NS-A and NS-B datasets where required. As only six out of 81 sequenced strains had the NS-A allele, only NS and NS-B datasets were used in the analyses.

We then subdivided the complete datasets of each gene according to viral traits, namely:

- host group (gull and duck)
- host type

- **BMG**: Black-headed Gulls (*Chroicocephalus ridibundus*) and Mediterranean Gulls (*Ichthyaetus melanocephalus*).
- **YAG**: Yellow-legged Gulls (*Larus michahellis*) and Armenian Gulls (*Larus armenicus*).
- **MD**: Mallards (*Anas platyrhynchos*).
- **OD**: Other ducks. This includes the common teal (*Anas crecca*), domestic duck (*Anas platyrhynchos domesticus*), garganey (*Anas querquedula*), northern shoveler (*Anas clypeata*), common coot (*Fulica atra*), and tufted duck (*Aythya fuligula*).
- HA subtype. Dataset was reduced to include subtypes H1, 2, 3,4, 5, 6, 7, 9,10, 11, 13 where greater than three sequences were available for statistical analyses.

#### Visualisation of phylogenetic incongruence

We inferred Maximum Likelihood (ML) phylogenetic trees for each gene segment using IQ-TREE, 1.5.5 (32) and ModelFinder (33) and obtained branch supports with SH-like approximate Likelihood Ratio Test (aLRT) and standard non-parametric bootstrap. All trees were rooted using the “best-fitting-root” function in Tempest v1.5 (34) and visualised in FigTree v1.4.2, with increasing node-order. To visualise incongruence, we traced the phylogenetic position of each sequence, coloured according to host, across unrooted ML trees for all internal gene segments. Figures were generated by modifying scripts from a similar analysis (35).

#### Quantification of nucleotide diversity

Complete alignments of each internal gene, as well as alignment subsets by host group, host type and HA subtype were used in “PopGenome” package in R v3.2 (36) to estimate nucleotide diversity. Per-site diversity was calculated by dividing the nucleotide diversity output by number of sites present in each alignment. As each subset contained different numbers of sequences, this value was normalised by dividing by the number of sequences in each respective dataset. Heat maps from this data were plotted in R v3.2.

#### Correlating traits with phylogeny (BaTS)

Null hypothesis of no association between phylogenetic ancestry and traits (host group, host type and HA subtype) was tested using Bayesian Tip-association Significance Testing (BaTS) beta build 2 (37) for all internal gene segments. Bayesian posterior sets of trees were inferred using MrBayes v3.2.6 (38) using the same segment-wise alignments generated for ML tree estimation. A set of scripts and commands used to generate the input file for BaTS are provided in [Supplementary materials]. Ratio of clustering by each trait on the gene segment trees that is expected by chance alone (Null mean), with the association that is observed in the data (Observed mean) was calculated. These expected/observed ratios were summarized in a heat-map with the y-axis ordered by the amount of reassortment observed. Data manipulation and figure preparation was done in R v3.2.

#### Quantification of diversity and between host transmission

Alignments generated for ML trees were also used in Bayesian phylodynamic analyses using BEAST v1.8.4 (39). We employed a strict molecular clock, a coalescent constant tree prior and the SRD06 site model with two partitions for codon positions (1^st^+2^nd^ positions, 3^rd^ position), with base frequencies unlinked across all codon positions. The MCMC chain was run twice for 100 million iterations, with sub-sampling every 10,000 iterations. All parameters reached convergence, as assessed visually using Tracer (v.1.6.0). Log combiner (v1.8.4) was used to remove initial 10% of the chain as burn-in, and merge log and trees files output from the two MCMC runs. Maximum clade credibility (MCC) trees were summarized using TreeAnnotator (v.1.8.4). After removal of burn-in, the trees were analysed using PACT (Posterior analysis of coalescent trees) (https://github.com/trvrb/PACT.git) to determine measures of diversity, and migration rates between hosts over time.

#### Geographical context for ‘Georgian origin’ internal protein coding gene segments

Internal gene sequences from, avian hosts, sampled across the world between 2005 and 2017 were obtained from gisaid.org (downloaded November 2017). Sequences (each segment separately) were divided into regions namely Asia (including Oceania), Europe, Africa, North America and South America. The program cd-hit-est (40, 41) was used to down-sample each regional dataset to 0.9 similarity cut-off level. These down-sampled sequences were then merged with the Georgian dataset. Discrete trait ancestral reconstruction with symmetric and asymmetric models were implemented in BEAST v1.8.4 (39) together with marginal likelihood estimation using path-sampling/stepping-stone analysis. The symmetric model was chosen over the asymmetric (log Bayes factor =14). The MCMC chain was run twice for 100 million iterations, with sub-sampling every 10,000 iterations. All parameters reached convergence, as assessed visually using Tracer (v.1.6.0). Log combiner (v1.8.4) was used to remove initial 10% of the chain as burn-in, and merge log and trees files output from the two MCMC runs. Maximum clade credibility (MCC) trees were summarized using TreeAnnotator (v.1.8.4). PACT was used to extract overall migration rates between trait locations.

## Results

### HA-NA subtype diversity and host-specificity

Over the six-year period between 2010 and 2016, 24 HA/NA subtypes of influenza A virus, including 12 different HA subtypes (H1, 2, 3, 4, 5, 6, 7, 9, 10, 11, 13, and 16) were isolated (Figure 1A). The diversity of subtypes varied from year to year, and associated with the level of prevalence in duck versus gull hosts. Within our sampling in Georgia, H9, H13 subtypes are found exclusively in gulls, while H1, H5, and H7 were detected exclusively in mallards. H3, H4, H6, and H10 were found in mallards and various other ducks. Positive evidence for multiple-species infection (ducks and gulls) was found only for H2 and H11 viruses in this dataset even though globally, many other subtypes are found in multiple hosts.

**Figure 1.**
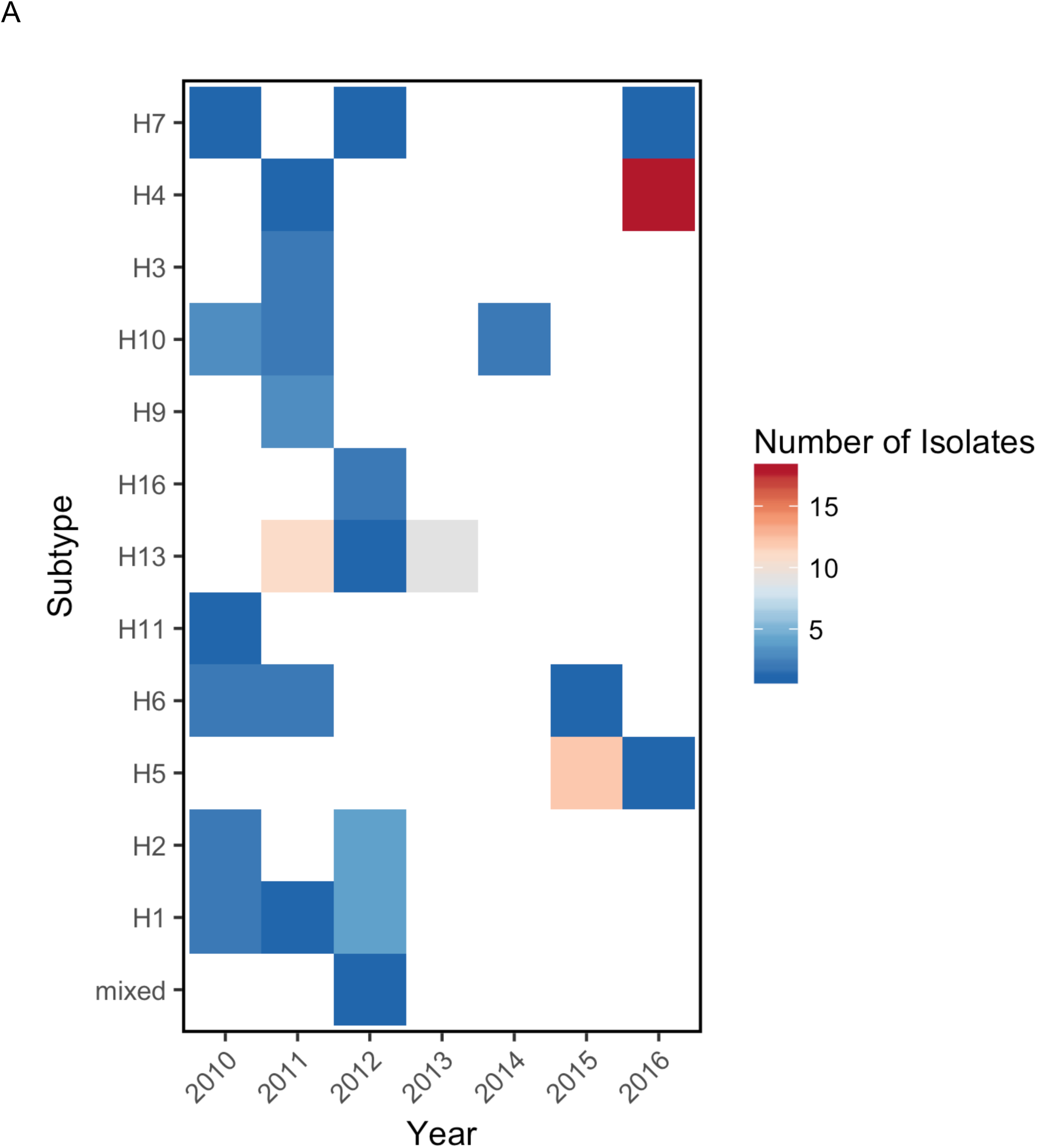
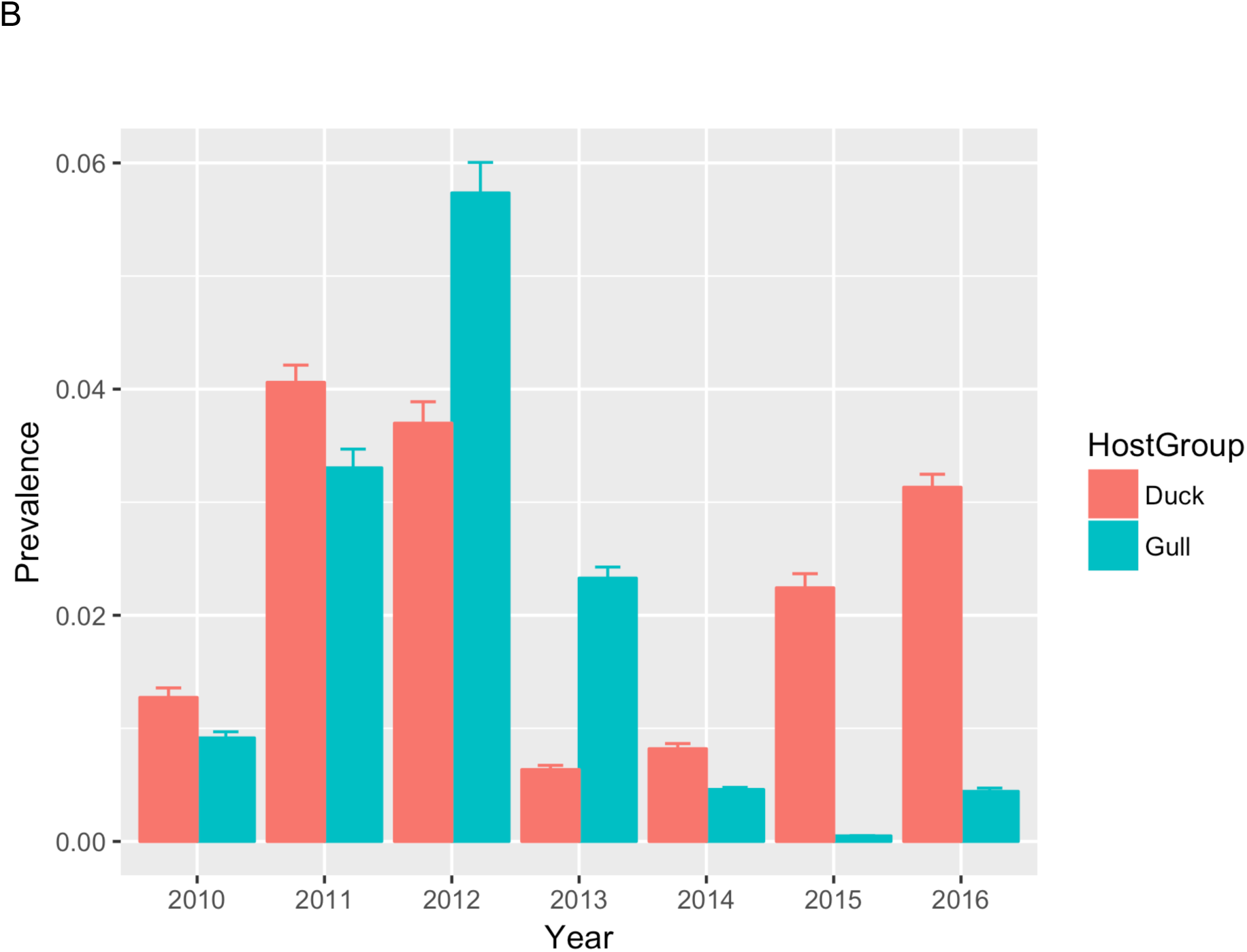
HA subtype-wise breakdown (A) and overall (B) yearly prevalence of viruses in Georgia during 2010-16. X-axis marks year of isolation. In bar chart 1A the Y-axis marks the proportion virus-positive samples +/− standard deviation and bars are colored according to host from which virus was isolated (duck in pink and gull in green). In heat map 1B, the Y-axis shows the HA subtypes of viruses isolated and squares are colored according to the number of isolates of each type identified.

In the first three years between 2010-12, up to seven different HA subtypes were found every year. These included H1, 2, 3, 4, 6, 10, 11, 13, and 16. H13, which was found in the greatest proportion of sequenced samples in 2011 and 2012 and was the sole type sequenced in 2013. In 2014, again only a single subtype was found (H10). The absence of more subtypes in these years could be explained by the comparatively low prevalence of IAV in these years, in both gulls and ducks in 2014 and especially ducks in 2013 (Figure 1B). In 2015, where prevalence was nearly zero in gulls, we saw HPAI H5 type viruses detected along with an H6. H4, which was previously isolated only in 2011, was the predominant type in 2016, followed by H5 and H7.

### Genetic structure of AIV detected in Georgia in 2010-16

For all gene segments except PA, there were two major subdivisions in tree topology – one clade containing sequences predominantly from ducks and one clade entirely derived from gull sequences (Figure 2, S2). The internal protein coding gene segments from certain subtypes formed sub-clades that were defined by year of circulation suggesting single-variant epidemic-like transmission within the population. This was seen in H13N8 in gulls and H4N6 and H5N8 in ducks. There were several examples of gull-derived viruses, which had several internal gene segments (other than NP) located in the ‘duck’ clade, mostly derived from Black-headed and Mediterranean Gulls (BMG). Only the PA gene phylogeny had an occurrence of a small sub-clade of Yellow-legged and Armenian Gull-derived (YAG) viruses clustered within the duck-derived viruses. For M gene segment, there were two major clades entirely defined by host species (except for 2 BMG viruses), and an outlier sub-clade consisting of H2 and H9 gull lineage viruses from BMGs. In PB1, PB2 and PA, these outlier-sub-clade viruses were found in various configurations in the tree. For NS, the tree topology divided into two alleles as reported previously (42). However, there were only six viruses from Allele *A* isolated from four mallards (MD), a garganey (OD) and a common teal (OD). Allele *B* splits into two sub-clades again defined by whether the viruses were isolated from gulls or ducks. The ‘duck’ sub-clade includes the outlier BMG viruses identified above for M. The long branch length to the gull sub-clade from the duck sub-clade in Allele *B* would suggest that there might be host-specificity in NS evolution, perhaps in response to differences between avian host innate immune responses.

**Figure 2.**
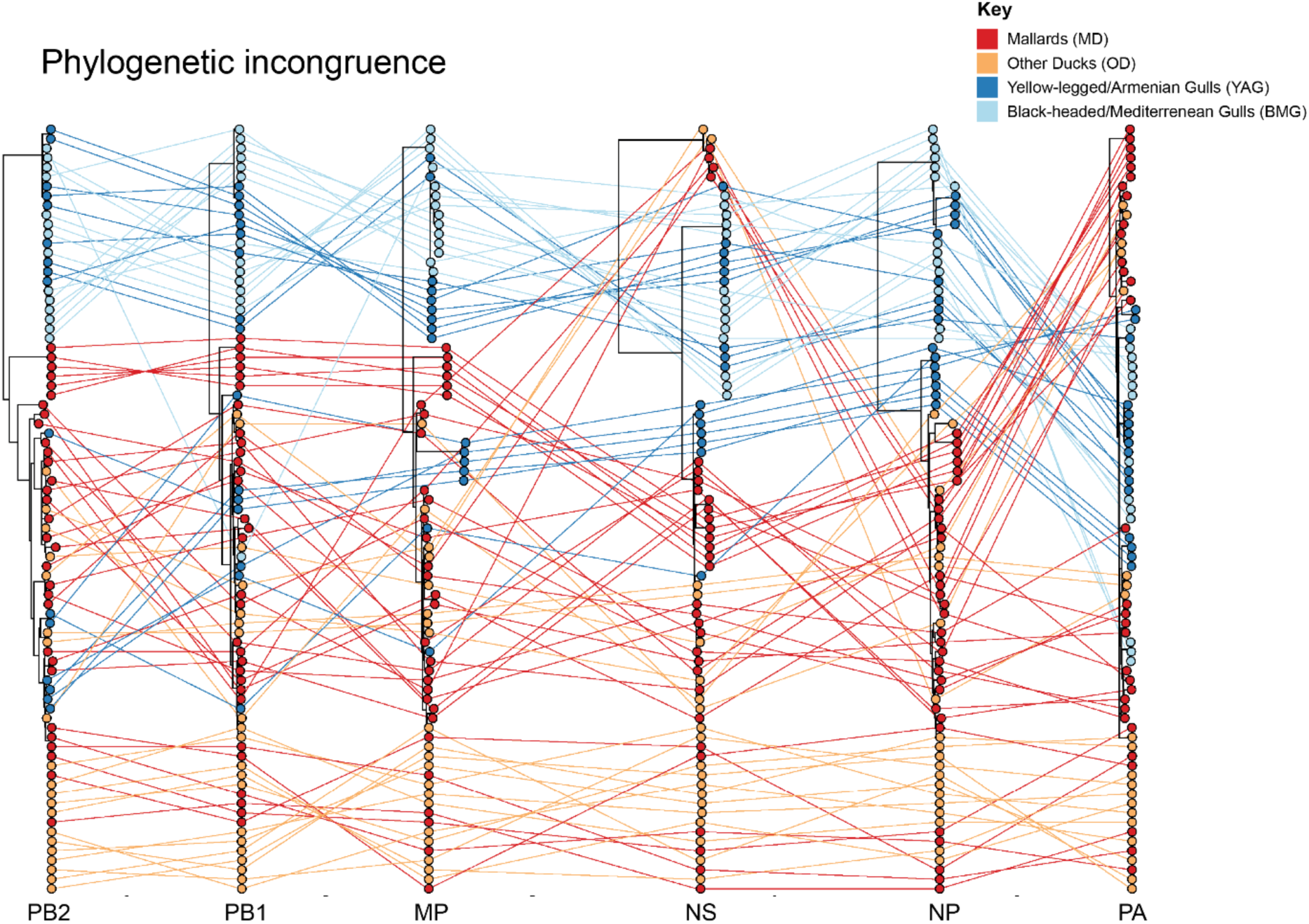
Maximum-likelihood trees for all internal genes – PB2, PB1, MP, NS, NP and PA, from equivalent strains connected across the trees. Tips and connecting lines are coloured according to host type –mallards (MD) bright red and other ducks (OD) orange, yellow-legged and armenian gulls (YAG) in blue and black-headed and mediterrenean gulls (BMG) in light blue).

### Variation in nucleotide diversity

We used the PopGenome package in R to calculate the per-site nucleotide diversity for all internal gene segments (Figure 3A-C). Nucleotide diversity of the internal gene segments in one surveillance site may be an indication of the breadth of sources where the viruses have been derived from. We found greater diversity in both gulls and ducks in gene segment NS (possibly because of the presence of both A and B alleles of this gene in the dataset) and PB2 (Figure 3A). When further sub-divided into “host types” as described in the methods, we found that the group of Black-headed and Mediterranean Gulls (BMG) had the highest per-site diversity. In comparison, the mallards (MD), the Yellow-legged and Armenian Gulls (YAG) and other ducks (OD) had relatively lower values across all internal gene segments, despite the OD comprising of a variety of ducks. Only the PA gene had greater diversity in Yellow-legged and Armenian Gulls than in Black-headed and Mediterranean Gulls (Figure 3B). When subset by HA subtype (Figure 3C), the internal gene segments associated with H4 and H13, the most abundant types found in our dataset, had the lowest diversity – possibly because several of the isolates were detected at the same time. Those less commonly isolated, such as H11 was detected in different years (2011, 2014) which may explain the high diversity of its NS, M, NP, PA, PB1, and PB2 gene segments. However, H3, which also has relatively high diversity were both detected at the same time (September 2011). Both NS and NS-B datasets were used in the analysis and as expected, the exclusion of sequences of NS-A (found exclusively in viruses from duck hosts), lowers the overall diversity within the ducks even when the values are normalised for the number of sequences found in each subset.

**Figure 3.**
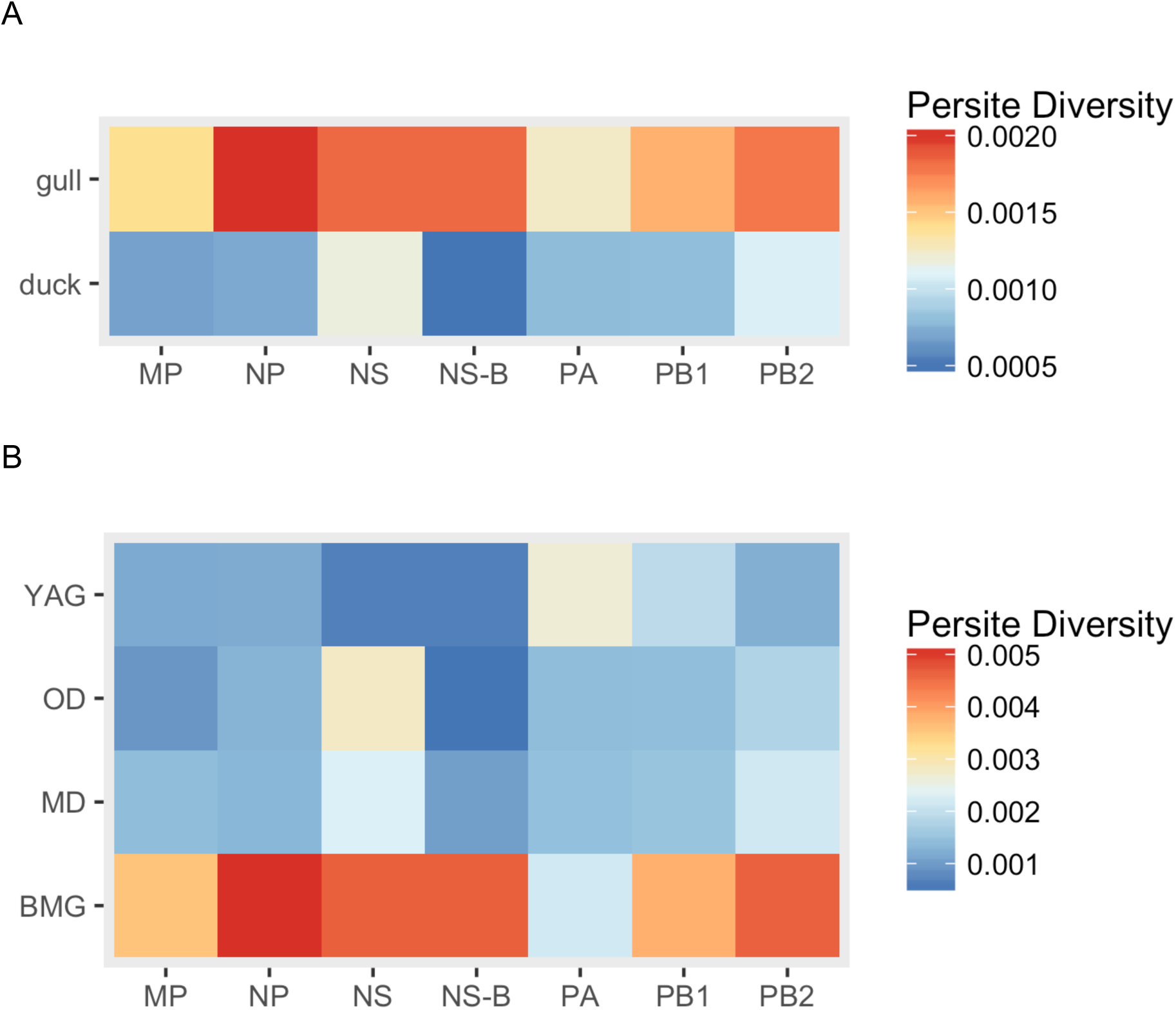
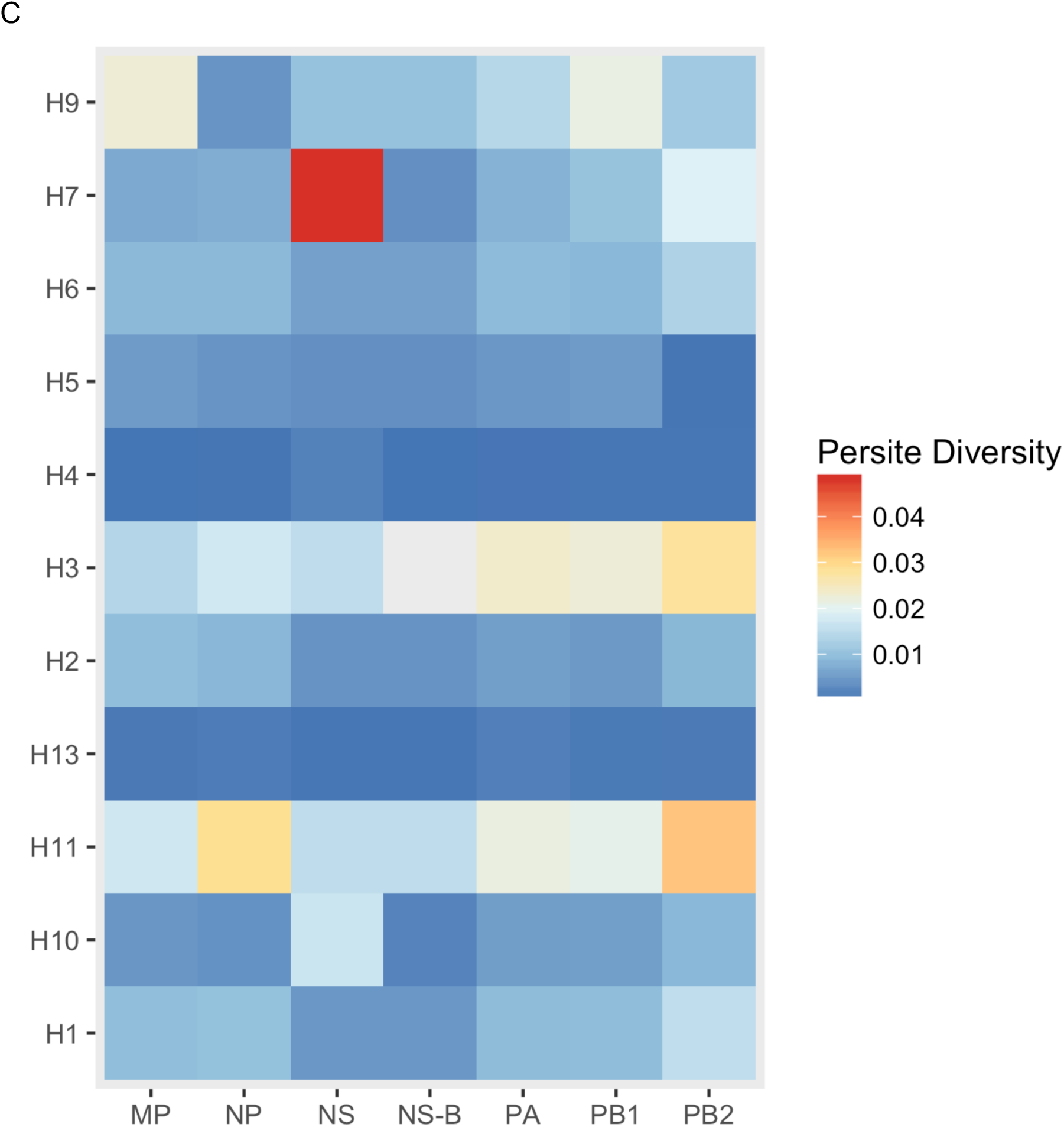
Overall per-site nucleotide diversity defined as average number of nucleotide differences per site between two sequences in all possible pairs in the sample population, normalised to the number of sequences in each population. Comparison between (A) gulls and ducks. (B) host-types (MD, OD, YAG, BMG) and (C) HA type are shown.

We tested the root-to-tip regression for ML trees for each of the six internal protein coding gene segments using Tempest v1.5 (34) to look for temporal signatures. All except NS gene showed positive correlation of distance with time, despite the short window of six years (Figure S1A). NS root to tip regression shows a negative slope, and it is likely confounded by the presence of two alleles A and B. Therefore, only NS-B allele, which forms a dominant portion of the NS gene segments in the data-set (75 out of 81), and shows clock-likeness (Figure S1B) were used for further analysis using BEAST v1.8.4. PACT analysis showed that the overall and yearly host-related diversity measures (Figure 4 A and B) show similar trends as seen in Figure 3.

**Figure 4.**
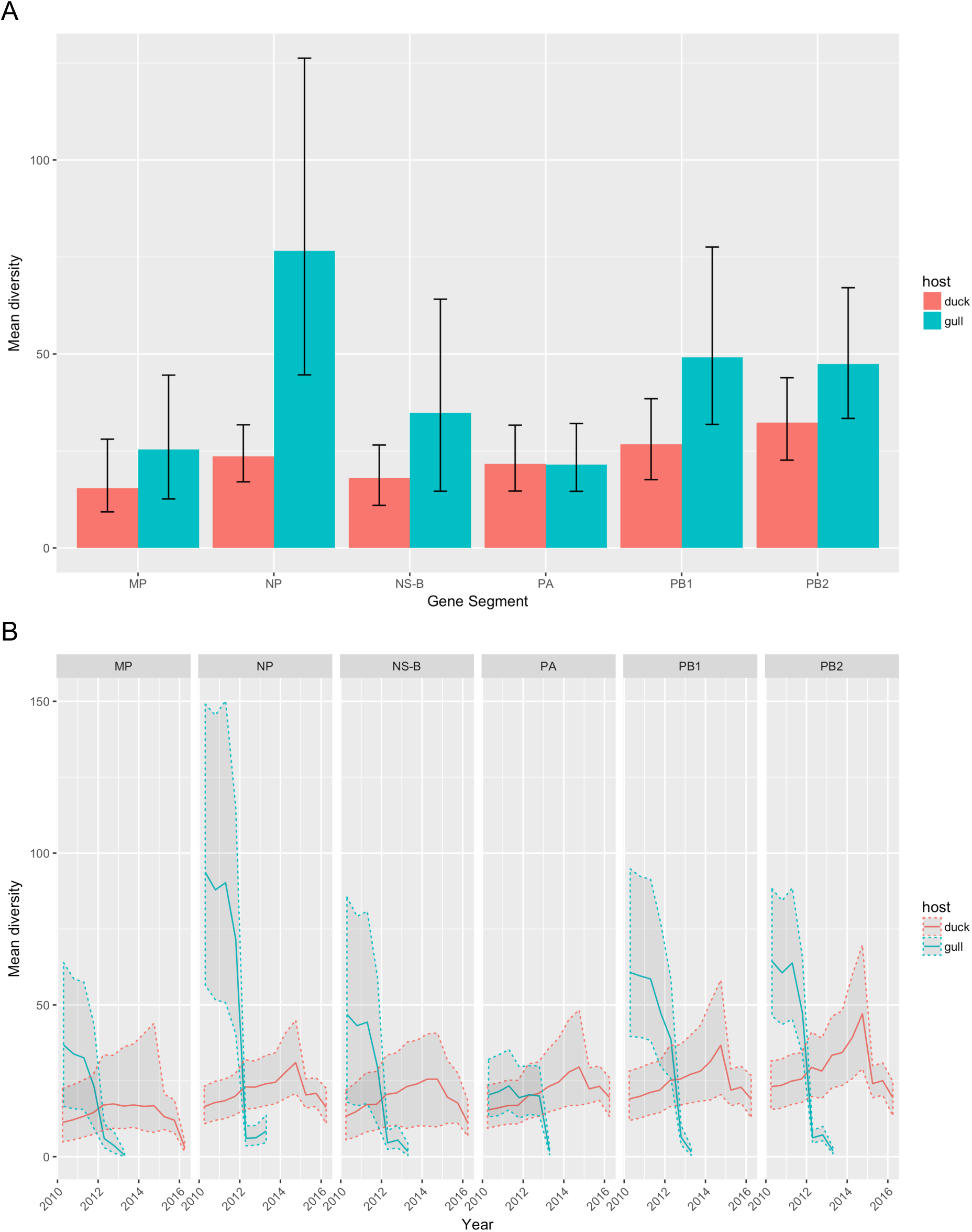
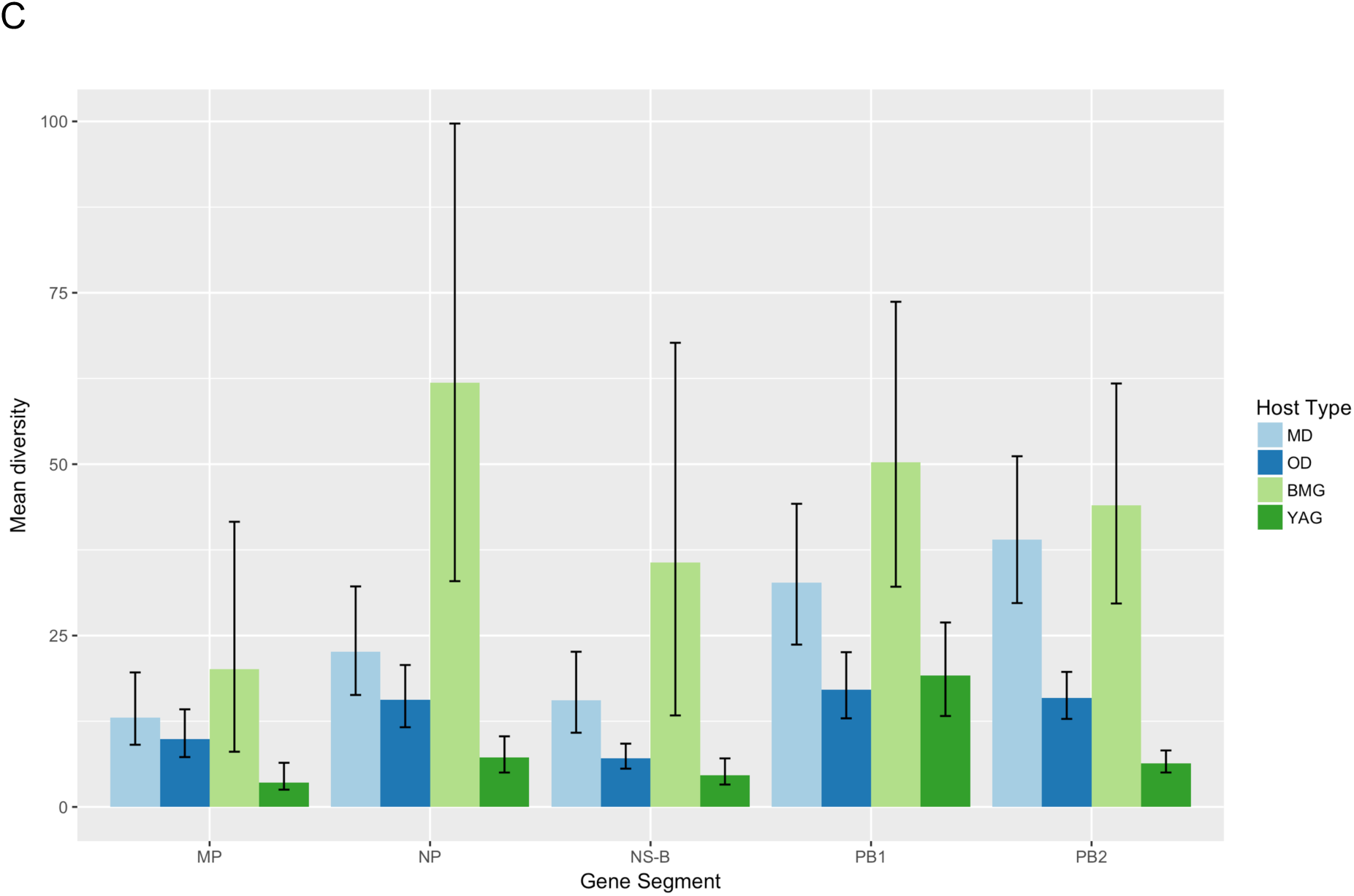
Overall/summary (A) and over-time/skyline (B) mean diversity for each segment from gulls (green) and ducks (pink) as determined by posterior analysis of coalescent trees (PACT). Here, diversity is defined as the average time to coalescence for pairs of lineages belonging to each host. Panel (C) shows overall/summary mean diversity values for ducks divided in to MD, OD (light and dark blue), and gulls divided into BMG and YAG (light and dark green).

### Correlation of traits with phylogeny

We tested the null hypothesis that there is no association between phylogenetic ancestry and traits (host group, host type and HA subtype) using Bayesian Tip-association Significance Testing (BaTS). Ratio of clustering by each trait on the gene segment trees that is expected by chance alone (Null mean), with the association that is observed in the data (Observed mean) are presented in Figure 5 (A-C). The higher the value of null/observed, the lower is the support for phylogenetic clustering of the given trait. Therefore, a higher value indicates a different ancestry. Hence, when we consider the HA subtype trait as “lineage”, it provides a measure of reassortment as described (43). Again, NS-B dataset was considered along with the complete NS dataset but no significant differences in trends were found. Panel A shows that gull viruses are more likely to cluster together in a phylogenetic tree than duck viruses in general. When viruses of gulls and ducks were further subdivided, panel B shows that OD viruses are less likely to cluster together in the tree, which is expected given that we have grouped together several duck species under this category. Among the rest, again it is the duck species (MD) that exhibit dynamic phylogenetic placing compared to both the gull types. The only exception is with the PB2 gene segment, for which the BMG show a lower level of phylogenetic clustering by species indicating putative reassortment events. When we consider the HA subtype (lineage) of the viruses, we find that H4 and H13, which showed the lowest nucleotide diversity, also show very low levels of reassortment, as does H5. There was not enough statistical power to interpret events in H1, 3, 6, 7, 9 or 11 viruses. Where statistically significant values were found, lower levels of clustering were observed.

**Figure 5.**
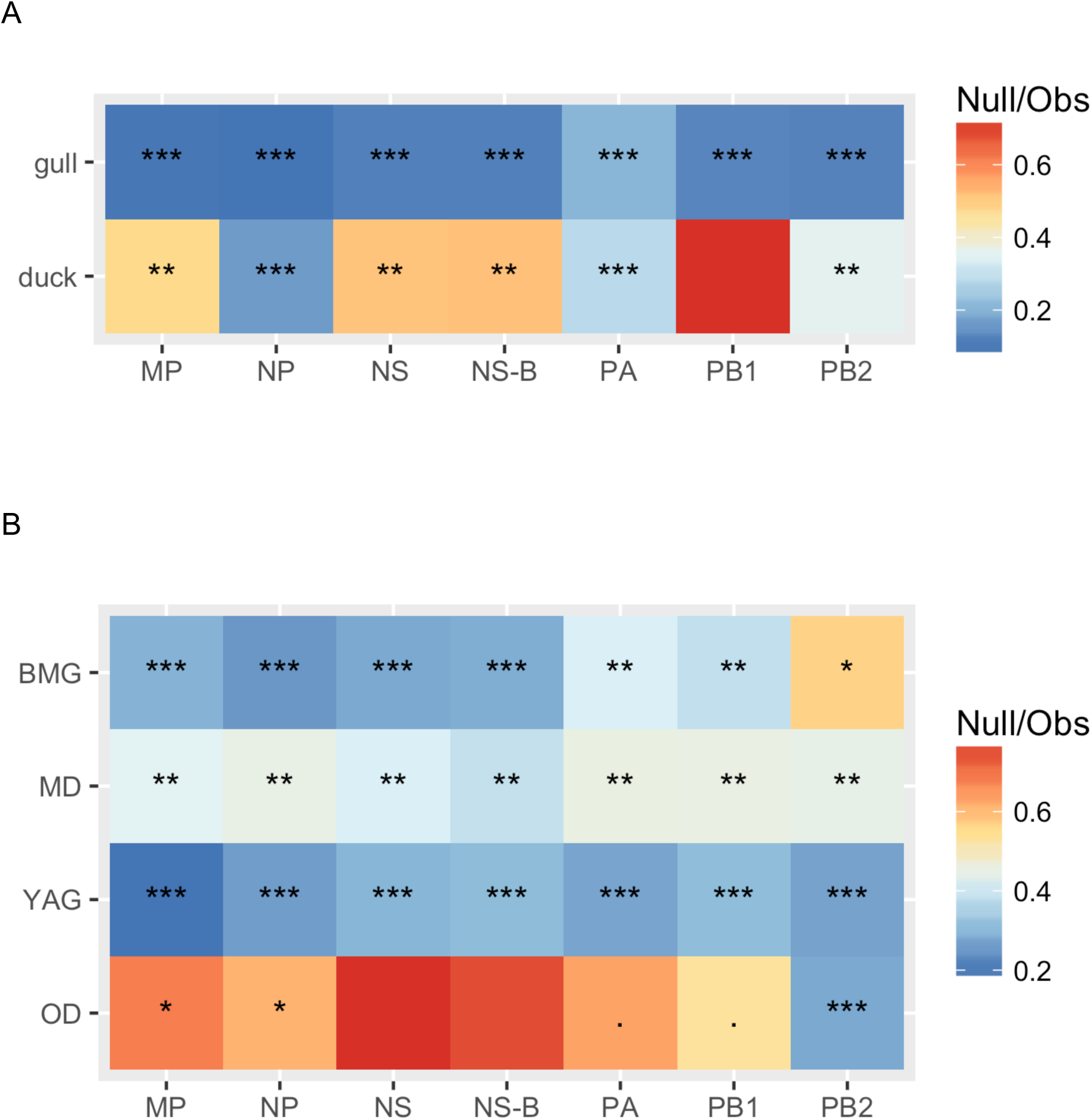
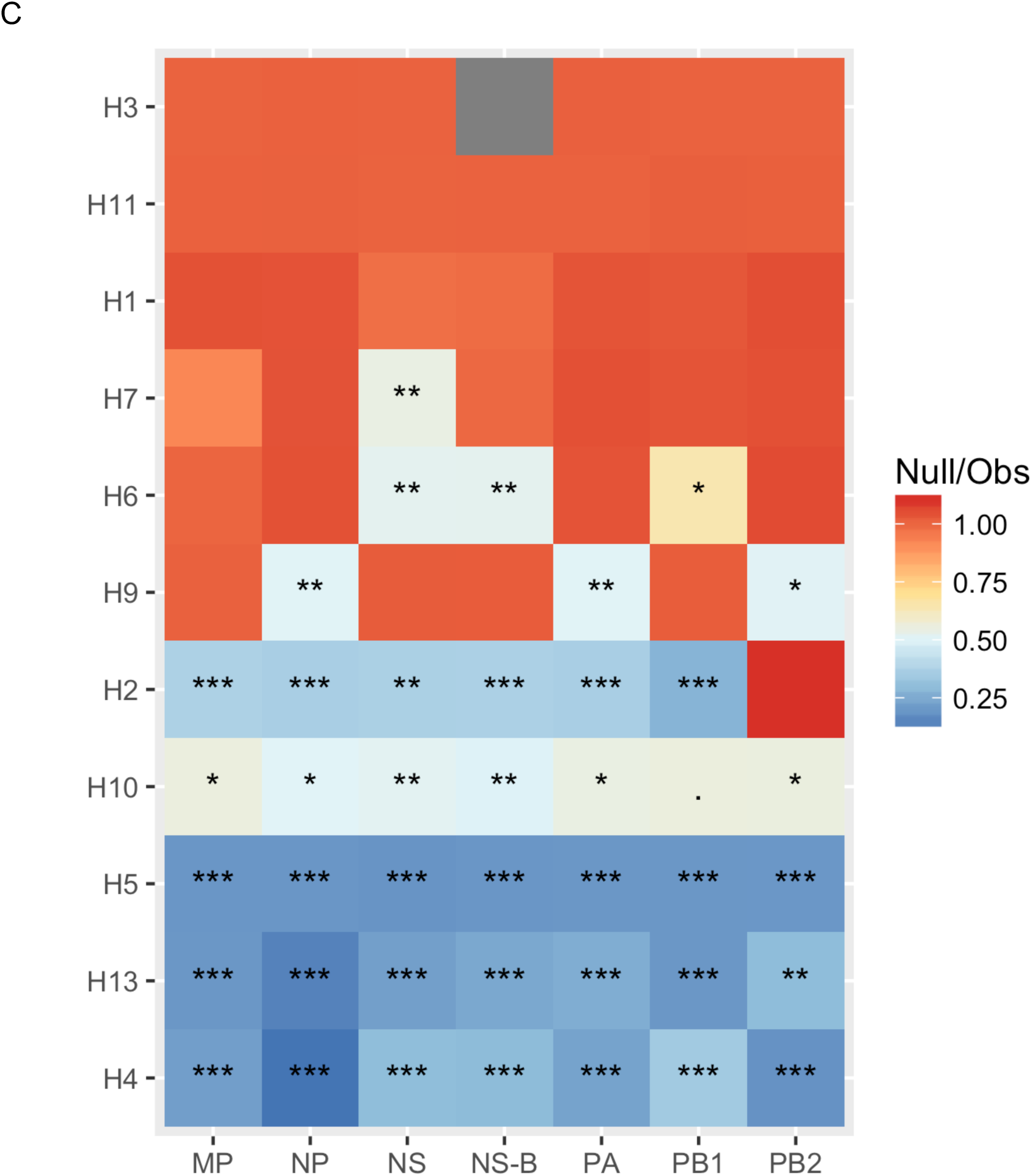
Summaries of expected/observed ratios from Bayesian Tip-association Significance testing (BaTS) for all internal genes. Higher values indicate less phylogenetic clustering by trait and hence higher rates of mixed ancestry. Comparison between (A) gulls and ducks. (B) host-types (MD, OD, YAG, BMG) and (C) HA type are shown. Asterisks indicate p-values (*** < 0.001, ** < 0.01, * < 0.05 and no asterisk > 0.05).

### Directionality of viral gene segment transfer

Figure S3 shows ancestral reconstruction of the host state along time-scaled phylogenies for five of six internal gene segments. The results are summarised in Figure 6A showing the mean number of host jump events from duck to gull and vice-versa. For all gene segments, most of the host spillover events are in the direction from ducks to gulls. In figure 6B we see that at a finer level, most of the host jump events happen within the duck (mallards (MD) to other ducks (OD)) and gull (Black-headed and Mediterranean Gulls (BMG) to Yellow-legged and Armenian Gulls (YAG) and *vice versa*) species. In transmissions from ducks to gulls it is largely noticeable only from MD to BMG. This likely explains the higher levels of nucleotide diversity and reassortment rates in the BMG viruses relative to YAG seen above.

**Figure 6A.**
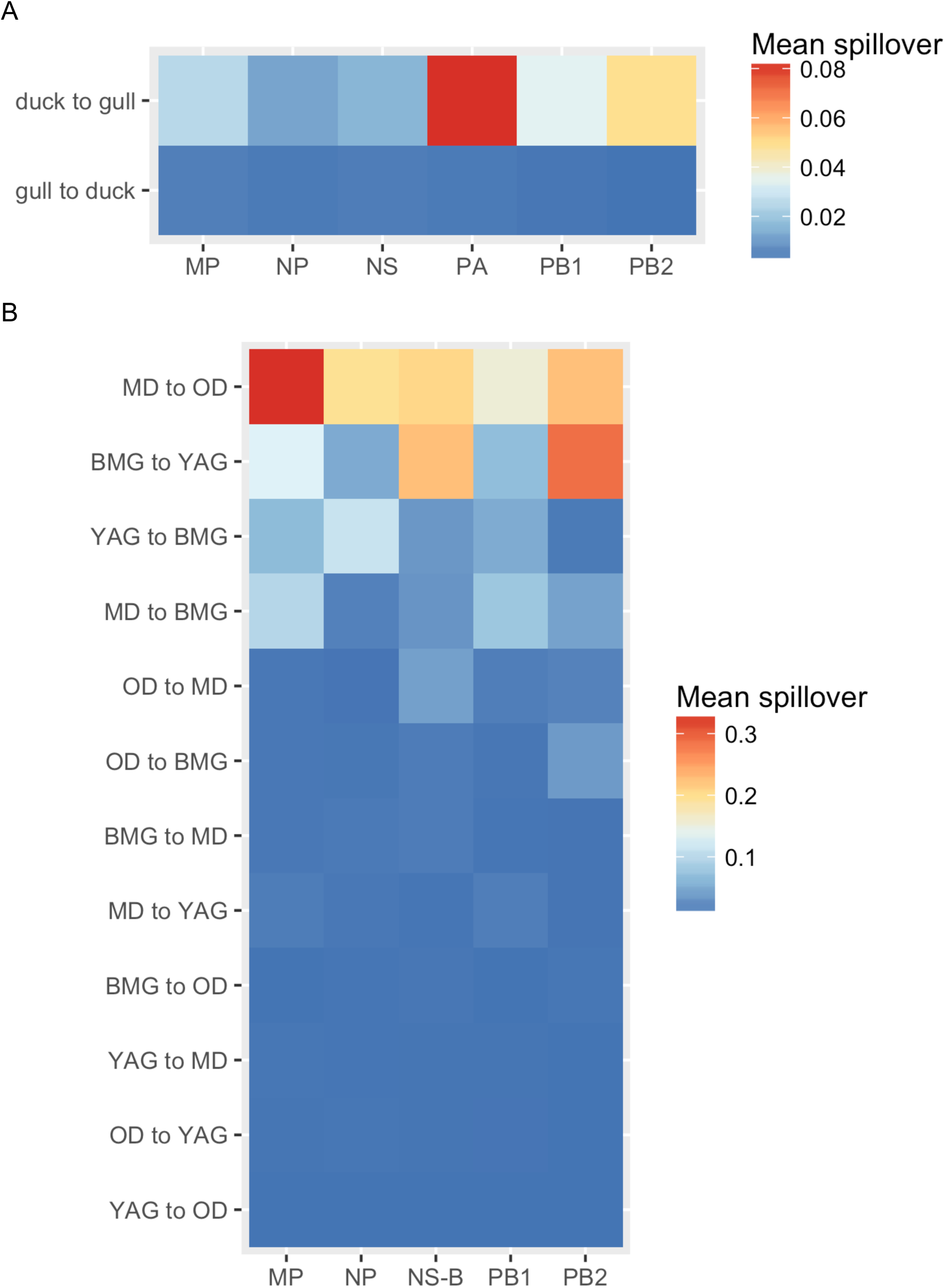
Summary of mean migration events between hosts in the direction from (A) duck to gull and gull to duck, and (B) between different host types - Mallards (MD), other ducks (OD), Black-headed and Mediterranean gulls (BMG) and Yellow-legged and Armenian gulls (YAG) derived from the genealogy.

### Geographical context for GE NS, M, NP, PA, PB1, PB2 segments

To determine the origin and destination of the internal protein coding gene segments found in viruses isolated in Georgia, we analysed our sequence dataset together with avian influenza sequences from a broader timeframe (2005-2016) and regional sampling. Figure 7A shows the genealogy for the NP gene for whose tips we know the location of sampling and whose internal nodes are estimated using discrete-state ancestral reconstruction in BEAST. Clades in which Georgian sequences occur are highlighted. Figure 7B summarises the genealogy in a circularised graph in which the arrowheads indicate the direction of transfer and the width of the arrow indicate the rate of transfer to different locations. The analyses reveal viruses from the Atlantic and Afro-Eurasian locations form largely separate clades, which is consistent with previous studies (44, 45). However, we do find instances of transmission across this divide, most notably to and from Asia and Europe. Many NP genes from Georgia cluster with other Georgian NP genes, in some cases forming the terminal branches spanning years indicating restriction to local spread. However, our dataset contains the latest Georgian sequences, and sequences from this timeframe were not available from the rest of Eurasia. Hence, we can expect to have missed identifying onward transmission. From the transmission we do identify, it appears that there is considerable migration into Africa and Europe and to a lesser extent to Southern/Eastern Asia. Most of the sequences transmitted into Georgia come from Asia and Europe, along with a single identified instance of direct transfer from North America.

**Figure 7A.**
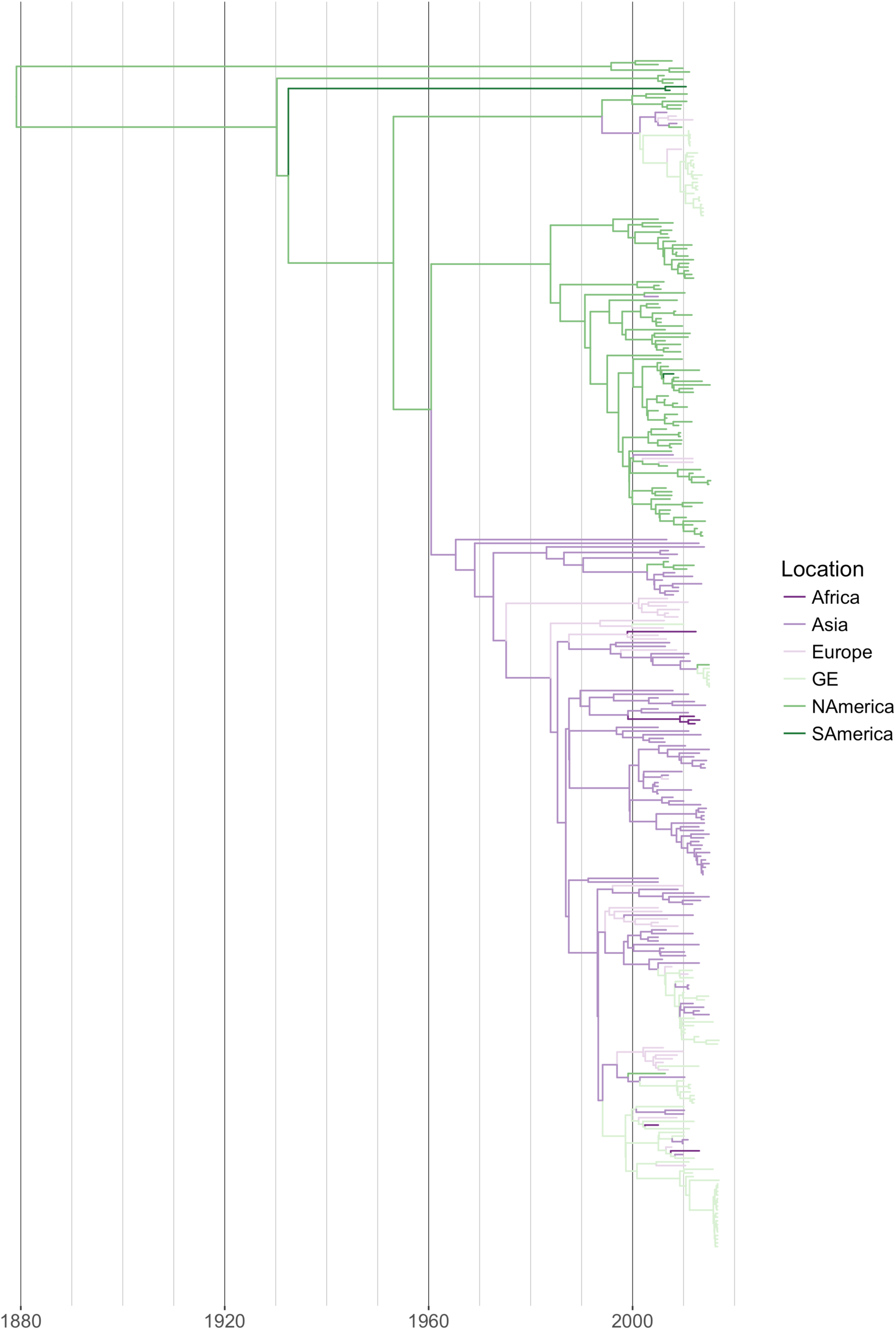
BEAST MCC (median-clade credibility) trees from viral sequences NP gene sequences isolated world-wide from avian hosts between 2005 and 2016. Branches are coloured according to location, observed at the tips and estimated at internal nodes by ancestral reconstruction of discrete trait. Africa, Asia, Europe in very dark, dark and light purple, Georgian sequences from this study in light green, North and South America in dark and very dark green.

**Figure 7B.**
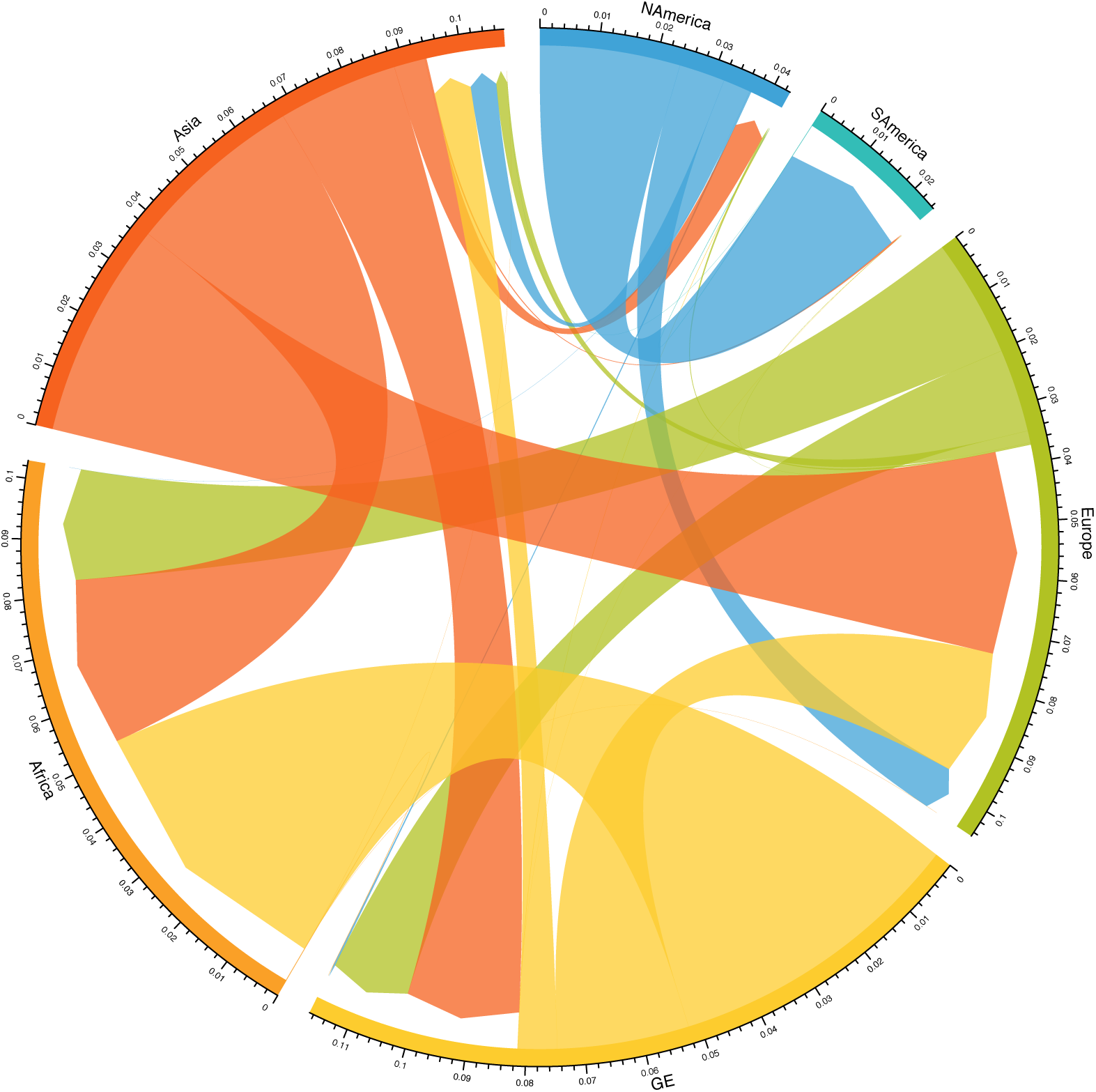
Circularised graph shows overall rates of migration, defined as the rate at which labels (locations) change over the course of the genealogy, between Georgia and other locations. Arrow heads indicate direction of migration; rates are measured as migration events per lineage per year (indicated by the width of the arrow). Asia in blood orange, Africa in orange, Georgia in yellow, Europe in green, South America in teal and North America in blue.

## Discussion

Wild birds have been shown to harbor substantial genetic diversity of avian influenza viruses. This study showed the diversity not only varied by year but was associated with the level of overall prevalence in different wild bird host species, perhaps influencing the observed rates and diversity if prevalence were low. From these results, there is little evidence that one species group maintains all influenza A virus diversity, there appears to be relative host-restriction in many subtypes (except for H2 and H11 viruses) and there are differences in prevalence dynamics depending on host. Therefore, one host is not representative of influenza A virus prevalence, dynamics and diversity across the wild bird reservoir. Within both ducks and gulls however, peak prevalence was consistently observed in hatch-year birds and with a more restricted subtype diversity, suggesting that there is an initial influenza A virus epidemic wave as naïve birds aggregate in their first year. Subsequently in the over-wintering period, a wider subtype diversity was observed in both host groups and adults were more frequently infected. This suggests that disease dynamics are complex and influenced by multiple host factors including age and annual life cycle stage.

It has previously been observed that some subtypes are routinely and nearly exclusively isolated from certain host families/genus, the most notable example being H13 and H16 viruses from gulls. However, mixed infections are relatively common but might be masked if subtype characterization requires virus isolation, therefore putting the clinical specimen through a culture bottleneck. Advances in sequencing direct from clinical material would more accurately (remove possible culture selection bias) establish the prevalence, subtype diversity and genetic diversity within wild birds.

In general, for all gene segments except PA, we identify strong patterns of clade topology defined by host. This suggests that there is segregated gene flow through these host populations with little inter-host reassortment. Additionally, within our study period there were large scale perturbations in ecology which might also influence our prevalence and subtype diversity estimates. For example, in 2014 and 2015 there was widespread reproductive failure in two gull host species due to nest flooding (Yellow-legged Gulls) and few returning adults to the colony (Armenian Gulls), and therefore few juveniles from which to detect the annual epidemic wave. The occurrence and significance of such ecological fluctuations on disease dynamics are unclear. We also increased the ability to sample migrant ducks in late summer and early autumn from August 2015 by constructing a duck trap in the newly created National Park. Again, this addition to sampling strategy likely increased the detection of influenza in these anseriform hosts as they were previously under-sampled.

We tested whether certain hosts maintained higher levels of nucleotide diversity in the non-immune related internal genes. PB2 and NS were the most genetically diverse in both gulls and ducks. Within host-group, Black-headed and Mediterranean Gull-derived viruses showed highest per-site diversity, Yellow-legged and Armenian Gulls lower diversity, likely because some of the viruses of the former were associated with reassortants probably derived from ducks (or another unsampled host group). While despite high rates of reassortment and spillover between duck subgroups mallards (MD) and other ducks (OD), the absence of any gull derived viruses in these ducks keeps their diversity levels lower compared to gulls/BMG.

Where gene flow does occur between host groups, for all gene segments, host-spillover events were in the direction of ducks to gulls and from other ducks to Black-headed and Mediterranean Gulls, likely explaining the higher levels of nucleotide diversity in these gulls observed above. Where HA and NA gene segments were acquired by gulls from ducks, there was a pre-requisite for a gull-clade internal gene cassette suggesting a host-restrictive effect for onward maintenance within the gull population (11, 46). Interestingly, Black-headed and Mediterranean Gulls only occur on the study site in the over-wintering period where there are also high densities of over-wintering ducks from other geographic areas. Although there is a duck-gull interface on the breeding grounds in summer, the duck densities are very much lower, perhaps suggesting that there is a threshold level of bird density that allows gene flow among hosts.

If we look at diversity by HA subtype, H4 and H13 were the least diverse and showed the lowest rates of reassortment and were also associated with hatch-year bird infections, suggesting a clonal expansion and epidemic gene flow through these birds. The 2014-2015 HPAI H5 epizootic also showed no reassortment unlike the 2016-2017 HPAI H5 viruses, perhaps indicating that the first wave of 2.3.4.4 viruses diffused through the wild bird population similarly to a ‘naïve’ infection, and subsequent epizootics have resulted in altered pathogen evolution strategies to maintain gene flow, similar to those previously observed in North America when considering the effect of latitude on gene flow (7).

When we examine the internal gene segments of the Georgian AIV in a broader geographical context, we find significant gene flow to and from Georgia with Europe and the rest of Asia, although data for Africa is very limited. Crossover into the Atlantic flyway appears to be mediated largely by gulls with some exceptions, notably the H5N1-NP gene that was transmitted between ducks.

From this study, the diffusion of avian influenza viruses within a multi-host ecosystem is heterogeneous. One host group cannot therefore be used as a surrogate for others. It is likely that virus evolution in these natural eco-systems is a complex mix of host-pathogen interface and ecological factors. Understanding such drivers is key to investigating these emerging pathogens, interpreting the data from different sites around the world and ultimately informing risk of incursion of emerging variants from one geographic region to another.

## Acknowledgements

This study including field work and sequencing was funded by National Institute of Allergy and Infectious Diseases, National Institutes of Health, Department of Health and Human Services contract No.HHSN2722000900007C and HHSN266200700010C “NIAID Centres of Excellence for Influenza Research and Surveillance” http://www.niaid.nih.gov/LabsAndResources/resources/ceirs/Pages/crip.aspx, and a DTRA FRCWMD Broad Agency Announcement (HDTRA1-09-14-FRCWMD GRANT11177182). The funders had no role in study design, data collection and analysis, decision to publish, or preparation of the manuscript. The sequencing data for this manuscript was generated while D. E. Wentworth was employed at the J. Craig Venter Institute. The opinions expressed in this article are the author’s own and do not reflect the view of the Centers for Disease Control and Prevention, the Department of Health and Human Services, or the United States government.

