## Supplementary Materials for "Avian influenza viruses in wild birds: virus evolution in a multi-host ecosystem"

**Supplementary figures**

**S1**. (A) Root to tip regression for ML trees generated from each internal gene of viruses isolated from Georgia 2010-16 using Tempest v1.5 and plotted in R v3.2. (B) Root to tip regression for ML trees generated from NS-allele B only sequences using Tempest v1.5 and plotted in R v3.2

A

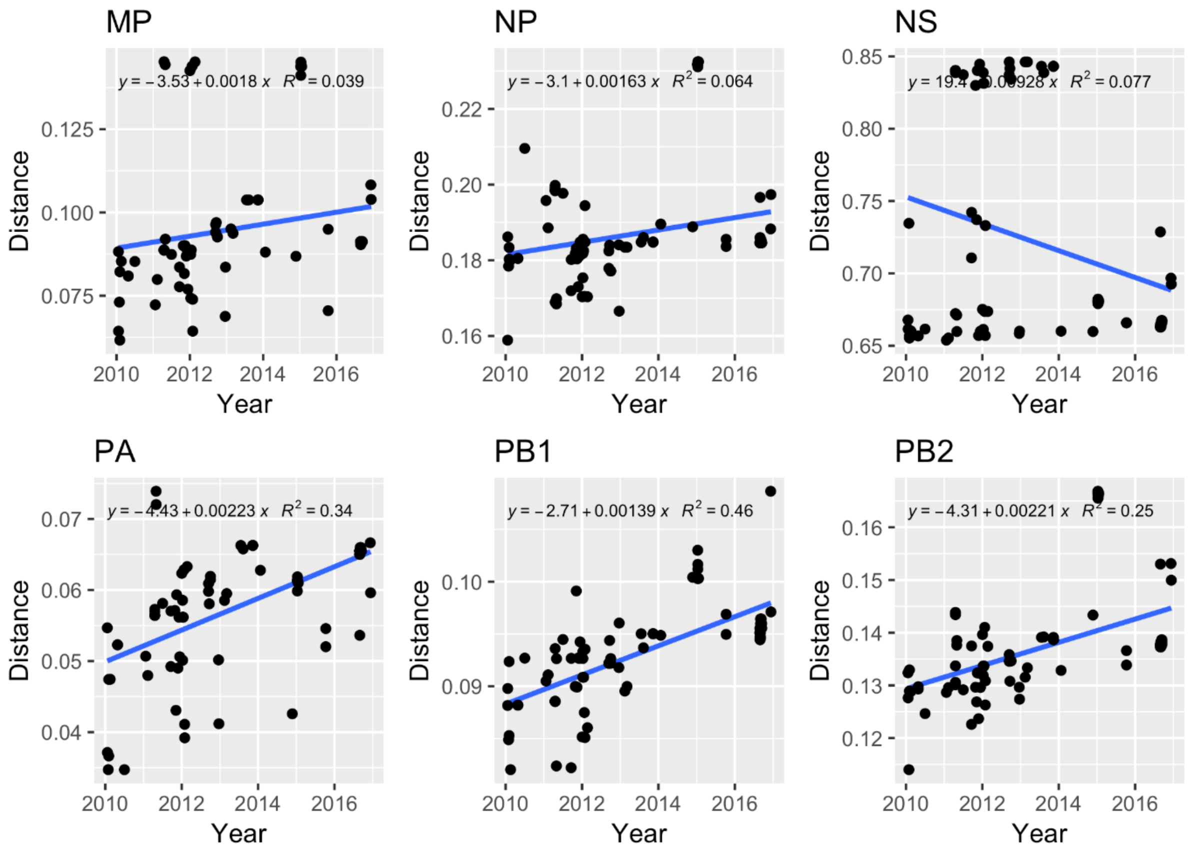


B

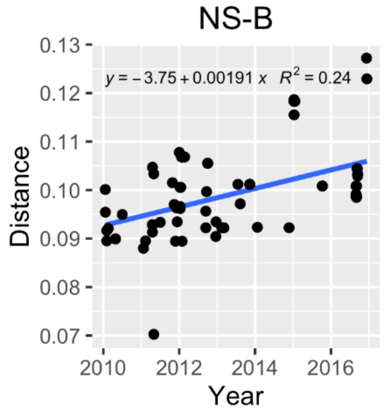

**S2.** Maximum-Likelihood trees for each gene segment of AIV isolated in Georgia 2010-16. Branch supports are indicated by the approximate Likelihood Ratio Test (aLRT) values. Tip labels are coloured according to the type of bird the strain was isolated from: Duck (pink), Gull (green).

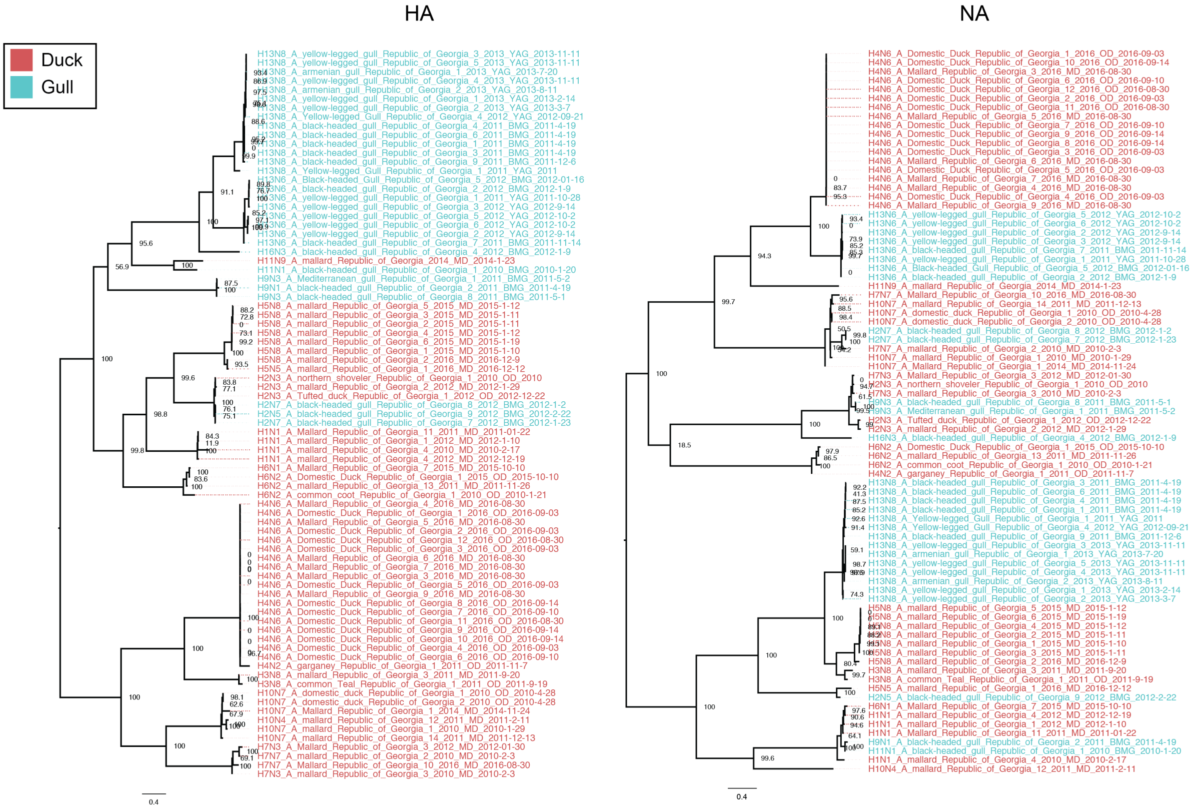

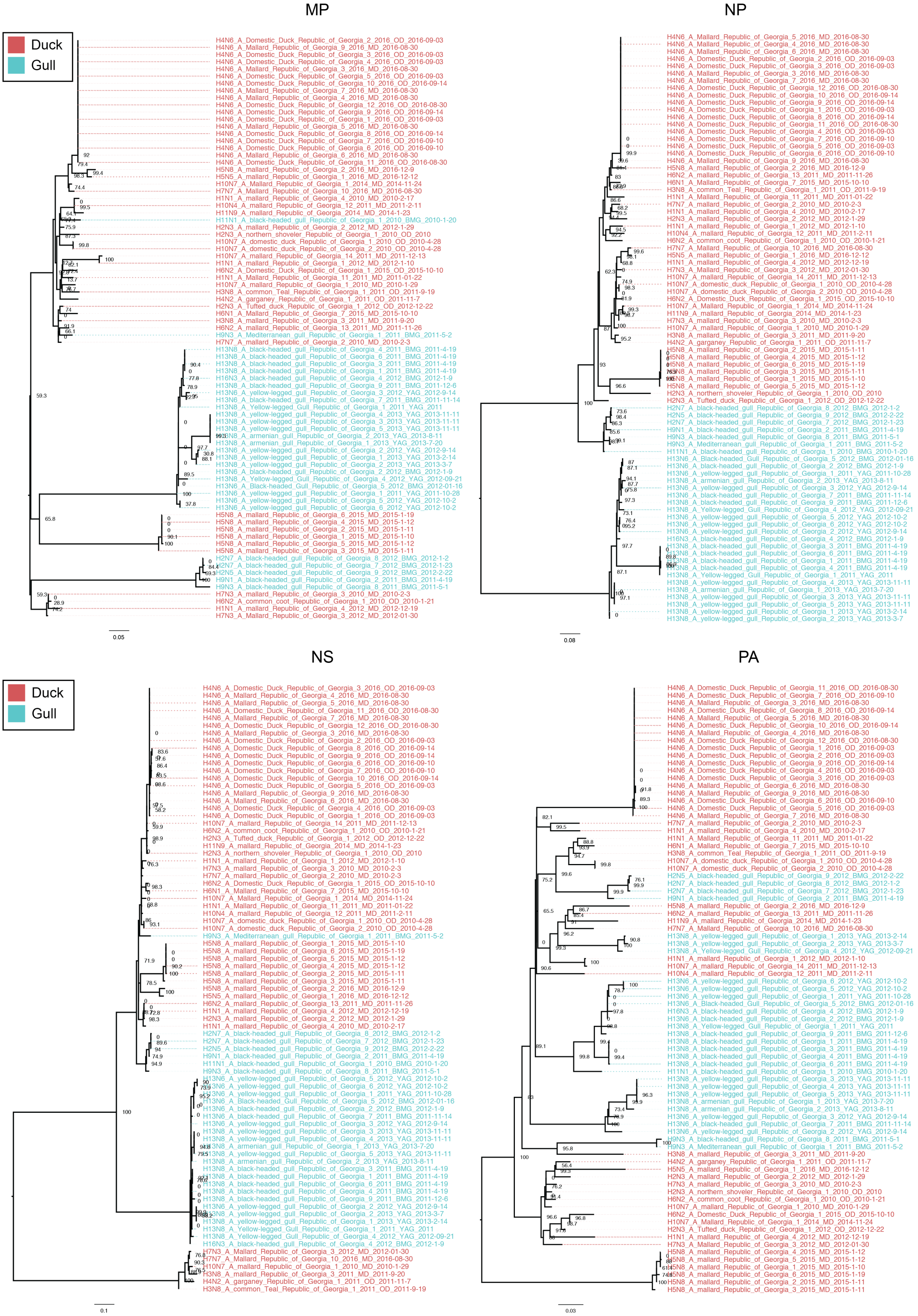

MP
NP
NS
PA
Duck
Gull
0.05
0.08
0.1
0.03

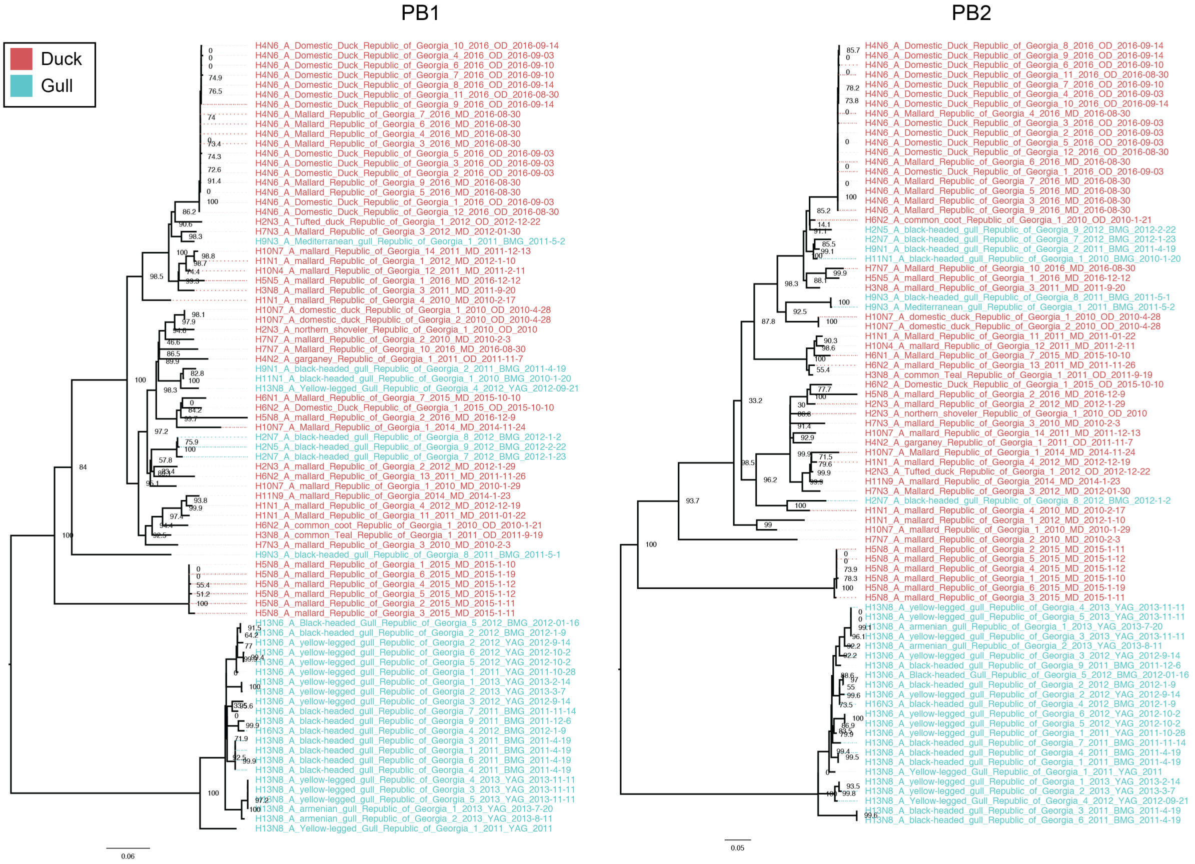


**S3.** Maximum clade credibility (MCC) trees for five of six internal gene segments of AIV isolated in Georgia 2010-16. Node icons are colored according to “host type” state inferred by BEAST v1.8.4. Black-headed and Mediterranean gulls (BMG)in red, Yellow-legged and Armenian gulls (YAG) in purple, Mallards (MD) in blue and other ducks (OD) in green.

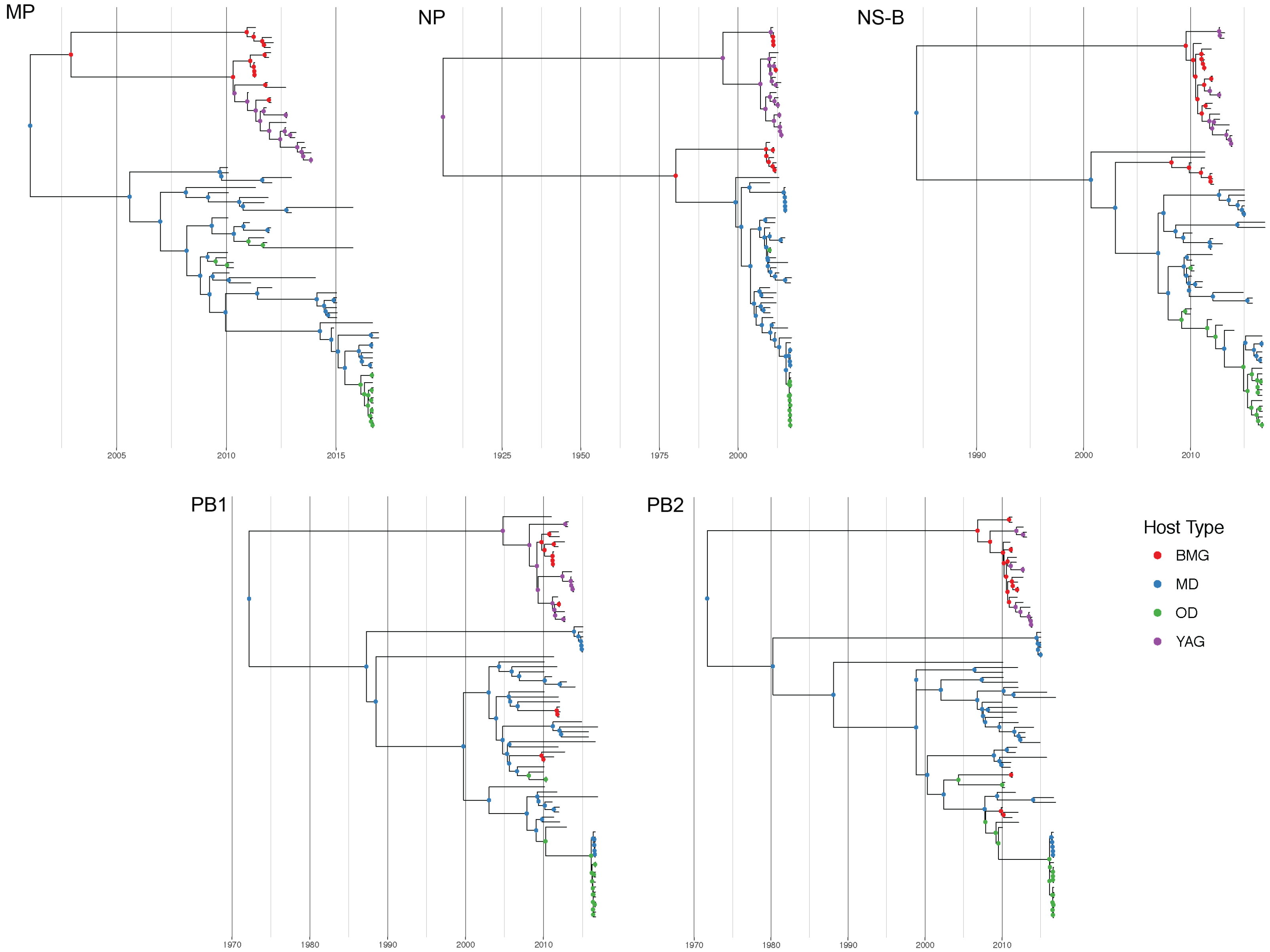
MP
2005
2010
2015
NP
1925
1950
1975
2000
NS-B
1990
2000
2010
PB1
1970
1980
1990
2000
2010
PB2
1970
1980
1990
2000
2010
Host Type
BMG
MD
OD
YAG
